# A dynamical-nonequilibrium molecular dynamics (D-NEMD) alanine scanning approach for identifying allosteric positions in proteins

**DOI:** 10.1101/2025.11.29.691197

**Authors:** Balazs Balega, Michael Beer, Dan T.S. Pike, James Spencer, A. Sofia F. Oliveira, Adrian J. Mulholland

## Abstract

Allosteric effects are widespread in proteins, but predicting the impact of sequence substitutions at positions distant from ligand-binding or enzyme active sites remains challenging, as their effects are mediated through complex dynamical networks. Pinpointing such positions experimentally is labour-intensive and low throughput. Equilibrium molecular dynamics (MD) simulations are useful: analysis methods can identify functional distal sites and networks, while free energy simulations can predict effects on stability or binding, given sufficient sampling; such simulations are typically time-consuming, requiring extensive simulation of each individual mutant. Here, we introduce a dynamical-nonequilibrium MD (D-NEMD) protocol using alanine substitutions as targeted perturbations to identify distal positions that influence enzyme function. Using the well characterised class A β-lactamase KPC-2 as a test system, we show that D-NEMD distinguishes functional from non-functional sites based on whether substitutions generate structured, long-range responses that reach the active site. KPC-2 is clinically important due to its broad substrate spectrum and the emergence of resistance-associated point mutations, many of which act through non-local effects on catalytic residues. Alanine substitutions at positions 179 and 164, which disrupt the Ω-loop salt bridge, trigger persistent structural responses propagating through the protein and reach both catalytic residues and the oxyanion-hole backbone. Likewise, alanine substitution at position 220 elicits pronounced responses extending to the active-site β-sheet and key loops, consistent with its known role in substrate specificity. In contrast, substitution at position 276—mutations of which have negligible kinetic impact—produces only local displacements with minimal propagation and no effect on catalytically relevant regions. These differential response patterns align with experimentally observed resistance phenotypes. D-NEMD therefore provides a fast, generalisable, and predictive approach for identifying allosterically connected distal positions. The protocol complements equilibrium MD, is straightforward to implement, and offers a tractable route for prioritising candidate sites for mechanistic study or future mutational scanning efforts.

**Statement of significance:** We introduce a dynamical-nonequilibrium MD (D-NEMD) protocol that uses alanine substitutions as the perturbation to identify residues, particularly those distal to an active or binding site, that modulate biological activity. We test this approach on the clinically important β-lactamase KPC-2: it distinguishes positions that elicit long-range responses that reach catalytic residues from another with only local effects, in agreement with experiment. The D-NEMD approach allows for tests of statistical robustness of responses to mutation, enabling prioritisation of candidate residues for experimental mutation. This should aid in identifying distal sites that may influence enzyme activity or serve other allosteric roles. The approach is generalisable to other proteins of biomedical and biotechnological interest.

## Introduction

Allostery—where the activity of a protein or enzyme is regulated by changes to the structure, dynamics, or binding state of a distant region—is a fundamental functional feature of many natural proteins. Allosteric communication can manifest through shifts in conformational ensembles, alterations in residue–residue interaction networks, or changes in local flexibility, and plays an essential role in modulating enzymatic activity, ligand binding, and protein–protein interactions. Despite substantial progress using experimental (e.g., mutagenesis, NMR, hydrogen–deuterium exchange) and computational approaches, reliably identifying which positions participate in functionally relevant allosteric coupling remains challenging, owing to the subtle and context-dependent nature of long-range communication.

Predicting the effects of sequence changes distal to the binding site in proteins therefore remains a major challenge of broad relevance in biomolecular science. Proteins are dynamic systems, and single-point mutations can alter stability, conformational flexibility, ligand or partner binding affinity, or catalytic activity in complex and often non-intuitive ways. Substitutions at positions far from an enzyme active site can nonetheless have substantial functional effects (1), frequently through changes in protein dynamics rather than direct structural disruption (1). Because these distal influences depend on nuanced, distributed conformational shifts, their outcomes are particularly difficult to anticipate. This challenge is especially relevant to β-lactamases, where single amino acid substitutions outside the active site can markedly alter the enzyme’s resistance profile (2–4).

β-lactamases are enzymes that hydrolyse β-lactam antibiotics, rendering them ineffective (5). β-lactam antibiotics are a cornerstone in the treatment of serious bacterial infections: they remain the most widely prescribed antibiotics. However, rising incidences of antimicrobial-resistant bacteria have proven one of the most pressing issues for global health in the modern world (6–8). The most common mechanism of β-lactam resistance in Gram-negative bacteria is the production of β-lactamases (5), which inactivate these antibiotics by targeting their cyclic amide group. Despite ongoing efforts to develop novel β-lactam drugs and β-lactamase inhibitors, the rapid and continuous evolution of bacterial β-lactamase variants presents a serious challenge, complicating both the effective prescription of existing β-lactams and the development of new drugs (9).

β-lactamases are grouped into four classes (classes A-D) (10) based on their sequence and structure, with class A being the most widely disseminated. Class A contains the *Klebsiella pneumoniae* carbapenemase (KPC) family, (11) with KPC-2 being the parent enzyme of >245 clinically observed variants as of January 2025 (12). This enzyme family is of particular interest due to its broad spectrum of activity: it has activity against all classes of clinically available β-lactam drugs, including carbapenems, considered to be “last resort” antibiotics (13). In addition to its broad substrate profile, KPC-2 has acquired point mutations that confer resistance to β-lactam/β-lactamase inhibitor drug combinations, such as ceftazidime/avibactam (14). It has been suggested that the high thermal stability of KPC-2 facilitates the accumulation of both point and insertion/deletion mutations, with some variants, such as KPC-40, KPC-64 and KPC-78, harbouring both types of changes (14).

KPC enzymes consist of two sub-domains, with the active site located in the cleft between them and a β-sheet (residues 234-237) forming the backbone (Figure 1a). The active site is surrounded by three key loop regions: the α3-α4 loop (residues 102-108), the Ω loop (residues 164-179), and the hinge region (residues 213-220). The Ω-loop contains two active site residues, E166 and N170 (16), both playing important roles in the activation and coordination of the deacylating water molecule. The deacylating water molecule is critical for catalysing β-lactam hydrolysis, as it attacks the acylated carbonyl group of the β-lactam ring within the acyl-enzyme complex, leading to the breakdown of the β-lactam ring and regeneration of the native enzyme (17). The stability of the Ω-loop, maintained by a salt bridge between R164 and D179 (Figure 1c), is thought to influence both the enzyme’s substrate profile and its catalytic turnover rate (18–20). Residues 234-243 form part of the β-sheet backbone (Figure 1b), with T235 and T237 playing crucial roles in substrate binding and carbapenemase activity (21). T237, in particular, forms part of the oxyanion hole with its backbone amide, along with the backbone amide of S70 (21). C238 forms a disulfide bond with C69, which in class A carbapenemases such as KPC has been shown to be essential for maintaining active site integrity (22–24). The hinge region includes T216 (Figure 1f), which interacts with the C-3 substituent of carbapenem substrates (25), and plays a key function in positioning the substrate for effective hydrolysis. Additionally, R220, also located in the hinge region, contributes to carbapenemase activity by forming a hydrogen bond network with E276 and T237 (26). Finally, the α3-α4 loop (Figure 1e) contains W105, a key active site residue for ligand recognition and implicated in resistance to β-lactamase inhibitors in KPC-2 (27).

**Figure 1:**
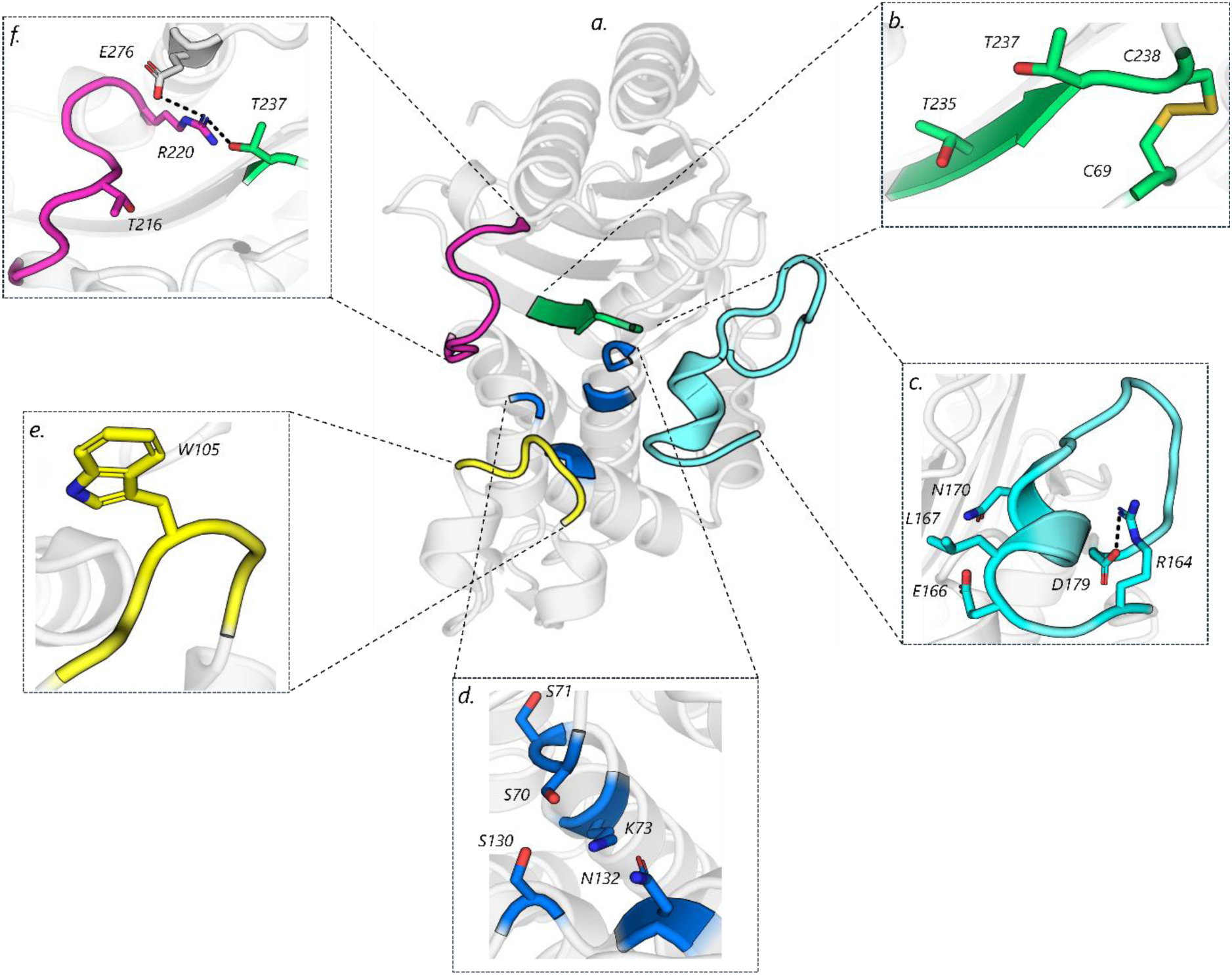
Structure of KPC-2 and important structural elements. ***a.*** X-ray structure of KPC-2 (PDB ID: 6D16 (15)), with functionally important regions highlighted in distinct colours: the Ω-loop coloured in cyan; the hinge region in purple; α3- α4 loop in yellow; active site β-sheet in green; and the core of the active site in blue. ***b.*** Close-up view of the β-sheet backbone in the active site, highlighting residues T235 and T237, and the disulfide bridge between C69 and C238. ***c.*** Structural detail of the Ω-loop, including residues E166 and N170, and the salt bridge between R164 and D179. ***d.*** Focused view of the core of the active site, highlighting the residues S70, S71, K73, S130 and N132 that line the catalytic pocket. ***e.*** Local view of the α3-α4 loop, including residue W105. ***f.*** Hinge region and residue T216, along with the hydrogen bonding network between R220, E276 and T237.

Point mutations in KPC-2, particularly in loop regions surrounding the active site, such as the Ω-loop (Figure 1c), have been shown to profoundly alter the drug resistance profile of enzyme variants by changing the dynamics of these structural elements (28). Understanding and predicting the (structural and dynamic) consequences of such mutations in KPC-2, and β-lactamases more broadly, should help in guiding effective use of β-lactam drugs and informing the design of novel antibiotics and β-lactamase inhibitors.

Extensive research has elucidated the (often complex) roles of key residues in KPC-2 activity, revealing how mutations at these sites can alter the resistance profile of enzyme variants (18, 28–31). However, and despite these substantial efforts, the ability to rapidly predict potential sites of emerging point mutations, and their subsequent functional effects, remains challenging. This difficulty is mainly due to the considerable time and resources required to pinpoint and characterise mutation sites across the entire protein using experimental methods such as site-directed mutagenesis (32).

To complement experimental efforts, computational approaches have increasingly been employed to elucidate the impact of key mutations, with equilibrium molecular dynamics (MD) simulations being the most commonly used method (33–37). While equilibrium simulations have provided valuable insights into the functional dynamics of β-lactamases, they are not suited for directly assessing how a specific local change gives rise to structured, protein-wide responses. Subtle mutation-induced effects can be difficult to distinguish from background conformational fluctuations, and extracting statistically robust signals often requires substantial simulation effort. (38–40).

To address these challenges, dynamical nonequilibrium MD (41–44) (D-NEMD) has seen growing application in the study of biomolecular systems (2,45–53). D-NEMD addresses limitations of equilibrium MD by applying a well-defined perturbation and quantifying the protein’s structural response relative to its equilibrium behaviour. By analysing the average response over a large ensemble of short trajectories, D-NEMD isolates reproducible, directional displacements that are specifically attributable to the perturbation, rather than to background thermal fluctuations. This capability makes D-NEMD particularly valuable for studying allostery, where the central question is how a local event—such as a mutation, ligand interaction, or conformational shift—affects distal functional regions of the protein.

In particular, for β-lactamases, previous D-NEMD simulations on KPC-2 and TEM-1, using ligand removal as the perturbation, have revealed the structural connection between active and allosteric sites. For both proteins, deletion of an allosteric ligand revealed residue networks linking known allosteric regions to the active site, highlighting the internal communication pathways within the enzymes (47). More recently, D-NEMD (using the removal of an active site ligand as the perturbation) identified a distal site that modulates KPC-2 activity (2), further demonstrating the method’s potential to uncover functionally relevant regions beyond the immediate catalytic site.

In this work, we present a novel D-NEMD application aimed at developing a rapid and efficient computational assay to probe the dynamical effects of point mutations in enzymes, using KPC-2 as a model system. This assay can be used to distinguish between sequence positions that influence catalytic activity and antibiotic resistance, and those with minimal functional impact under the tested conditions. Within this framework, the instantaneous substitution of a target residue with alanine serves as the external perturbation in D-NEMD, enabling the observation of how structural responses evolve and propagate throughout the enzyme. This approach facilitates the identification of functional connections between target residues and the active site, offering predictive insight into how mutations at these positions may alter active site dynamics and, therefore, the enzyme’s activity profile. To evaluate the assay’s ability to distinguish between functionally impactful sites and those with minimal effect, we simulated four KPC-2 positions: 164, 179, 220, and 276. Of these, three (164, 179, and 220) are experimentally known to affect enzyme activity (14, 18, 20, 26, 29, 31), while position 276 is considered functionally neutral (26). D-NEMD effectively differentiated between these sites, based on whether the structural changes induced by alanine substitution propagated to the enzyme’s active site.

## Materials and Methods

### Equilibrium MD Simulations

Equilibrium MD simulations of KPC-2 in complex with a heteroaryl phosphonate non-covalent inhibitor (PDB: 6D16 (15)) as described in Beer *et al.*, 2024 (2), were used as the starting structures for the nonequilibrium simulations presented here. In brief, these simulations were carried out with GROMACS 2019.1 (54), using the AMBER99SB-ILDN (55) forcefield to describe the protein and GAFF (56) parameters for the ligand. The system was solvated in TIP3P water (57) with 200 mM NaCl, equilibrated under NVT and NPT conditions, and simulated for 250 ns across five independent replicates. Temperature and pressure in the NPT production runs were maintained constant using a first-order stochastic velocity rescaling thermostat (58) and the Parrinello-Rahman pressure barostat (59). System equilibration was checked by analysing the Cα RMSD (Figure S2), with convergence below 2 Å indicating that the global protein conformation had relaxed and remained stable over the production timescale.

### D-NEMD simulations

Frames were extracted every 5 ns from the equilibrium trajectories described above, spanning the interval from 50 ns to 245 ns, resulting in a total of 200 frames (Figure S1). In each frame, the residue at the position of interest (specifically R164, D179, R220, and E276) was instantaneously mutated to alanine, with the new system subsequently simulated for 1 ns using the same settings and parameters as the equilibrium simulations reported by Beer *et al*. (2). This procedure yielded four sets of 200 nonequilibrium simulations, each targeting a different KPC-2 residue, namely R164, D179, R220, and E276. Of these, positions 164, 179, and 220 are considered functionally relevant, with experimental evidence indicating that mutations at these sites affect enzymatic activity (14, 18, 20, 26, 29, 31). In contrast, position 276 was selected as a non-functional control, as mutations at this site have been experimentally shown not to impact activity (26). The response of the protein to the alanine mutations introduced was then extracted using the Kubo-Onsager relation (60), as described in detail in Balega *et al.* (61). From this analysis, the evolution of the average Cα displacement vector, between every pair of nonequilibrium and equilibrium trajectories, was calculated.

In addition to the nonequilibrium simulations involving alanine mutation, a set of control simulations (termed “null perturbation”) (2, 62) was carried out following the same workflow (Figure S1). This control randomises atomic velocities at the same intervals along the equilibrium trajectory, rather than applying a structural perturbation. The changes stemming from the null perturbation represent intrinsic thermal motions of the protein. Subtraction of these motions from the nonequilibrium responses at the same timepoints effectively removes noise. This analysis reveals the specific response to the mutation, thereby enabling precise mapping of its structural and dynamic impact. The average Cα displacement vectors shown in Figures S3-S6 (computed after removing thermal fluctuations via the “null perturbation” analysis) indicate the direction of the mutation-induced structural changes at each time point, while their norms quantify the corresponding response amplitude (Figure S7-S9). The evolution of both the direction and amplitude of the protein’s responses was visualised using PyMOL (63), providing a detailed description of how these structural changes propagate from the mutation site to the active site. This analysis enables the identification of internal structural connections between these regions and whether the mutation effects extend to functionally critical regions of KPC-2.

The statistical significance of the average responses was evaluated by computing the standard error of the mean for each component of the displacement vector, as described in Balega *et al.* (61). Only vector components with magnitude exceeding their estimated uncertainty are shown throughout, ensuring that the reported responses reflect genuine mutation-induced signals rather than thermal noise. Mutations at positions associated with altered resistance profiles, specifically R164, D179, and R220, triggered response pathways that affect catalytic regions (Figure S8).

## Results and Discussion

We use alanine substitutions as the external perturbation in D-NEMD. The aim is a computational assay to predict whether mutations at these positions influence enzyme activity through long-range structural communication. KPC-2, a well characterised class A β-lactamase and an important biomedical target, is used here as a test system to evaluate the ability of D-NEMD to differentiate between positions that significantly influence enzymatic activity and those with minimal functional impact.

An important target in the fight against antibiotic resistance, KPC-2 exhibits broad-spectrum activity; it is active against many β-lactam antibiotics (9). KPC-2 can acquire point mutations that change its spectrum of activity. Such mutations confer resistance to β-lactam antibiotic/inhibitor combinations such as ceftazidime-avibactam (14, 18, 20, 28–31). Four KPC-2 positions were selected here for alanine mutation, including three associated with altered resistance phenotypes (R164, D179, and R220) and one (E276) where mutation has been shown experimentally not to affect activity and is chosen as a control (26).

Positions 179 and 164, an aspartate and arginine, respectively, in KPC-2, are both located in the Ω-loop (Figure 1c) and play essential roles in the conformational stabilisation of this region through a salt bridge between the two residues (14). Mutations at these sites are known to disrupt this critical interaction. Several studies have shown that Ω-loop dynamics are important in the positioning of catalytic residues N170 and E166, which together are responsible for orienting and activating the deacylating water molecule (20, 64, 65). Consequently, KPC-2 variants harbouring point mutations at positions 179 (e.g. D179A, D179Y) and 164 (e.g. R164A, R164S) exhibit altered activity profiles compared to the wild-type enzyme (14, 20), probably due to the disruption of the D179-R164 salt bridge. Clinically observed and experimentally characterized mutations at these positions are associated with significant increases in minimum inhibitory concentrations (MIC) for ceftazidime, and reductions in carbapenemase activity, compared to KPC-2 (14, 29, 31).

The effect of alanine mutation at position 220 was also examined here. This site is in the hinge region; in KPC-2, it is an arginine residue known to play a pivotal role in the hydrolysis of carbapenems (26). R220 forms a hydrogen bond with T237, a threonine residue situated within the β-sheet backbone of the active site (Figure 1f) (16, 22, 26). This interaction is crucial for maintaining the correct orientation of T237, which in turn, forms hydrogen bonds with the C3 carboxylate group of carbapenem substrates, an arrangement essential for effective deacylation of carbapenem acyl-enzymes (26). Site-directed mutagenesis showed the importance of a positive charge at position 220 for substrate processing (16, 25, 26, 66): substitution of R220 with alanine (R220A) substantially decreases turnover number (*k_cat_*) for carbapenems (26). Conversely, this mutation also enhances hydrolysis of penicillin-class β-lactams, suggesting that R220 contributes to both substrate specificity and the catalytic mechanism of KPC-2 (26).

Previous work by Papp-Wallace *et al.,* identified that the KPC-2 (E276A) mutant had negligible effect on the catalytic activity of the enzyme against a range of β-lactam substrates, when compared to the parent enzyme (26) Hence, an alanine substitution in this position is used as a negative control, in order to demonstrate the usability of this technique in identifying residues that are highly important for function.

For each position, the effect of perturbation i.e. substitution by alanine was investigated by D-NEMD. Dynamical responses were calculated and analysed to assess whether these changes propagate to the enzyme’s active site. To ensure reliable and converged dynamical responses, 200 nonequilibrium simulations, each of 1 ns duration, were conducted for each site of interest. The choice of a 1 ns nonequilibrium trajectory length reflects the fact that mutation-induced structural responses in D-NEMD occur on very short timescales (61). For relatively small systems such as KPC-2, the dominant displacement patterns emerge as early as 10 picoseconds (Figure S7). This short (1 ns) timescale also minimises computational cost. 1 ns trajectories capture dynamical responses while keeping the computational requirements low enough to facilitate high-throughput analysis of many mutation sites.

### Response to Alanine Substitution at Position 179

As mentioned above, D179 forms a functionally important salt bridge with R164 (14). It also participates in an intra-Ω-loop hydrogen bonding network involving R161, D163, R164, S171, D176 and R178, with additional contacts to P67 and L68 in the loop preceding the catalytic core (20). In D-NEMD, substitution of D179 with alanine (D179A) disrupts these key interactions, triggering rapid and widespread structural responses across the enzyme (Figure 2). These changes, which originate in the mutation site, are transmitted to the active site, highlighting strong structural communication between the two regions that underpins the experimentally observed functional modulation (18, 20, 31).

**Figure 2:**
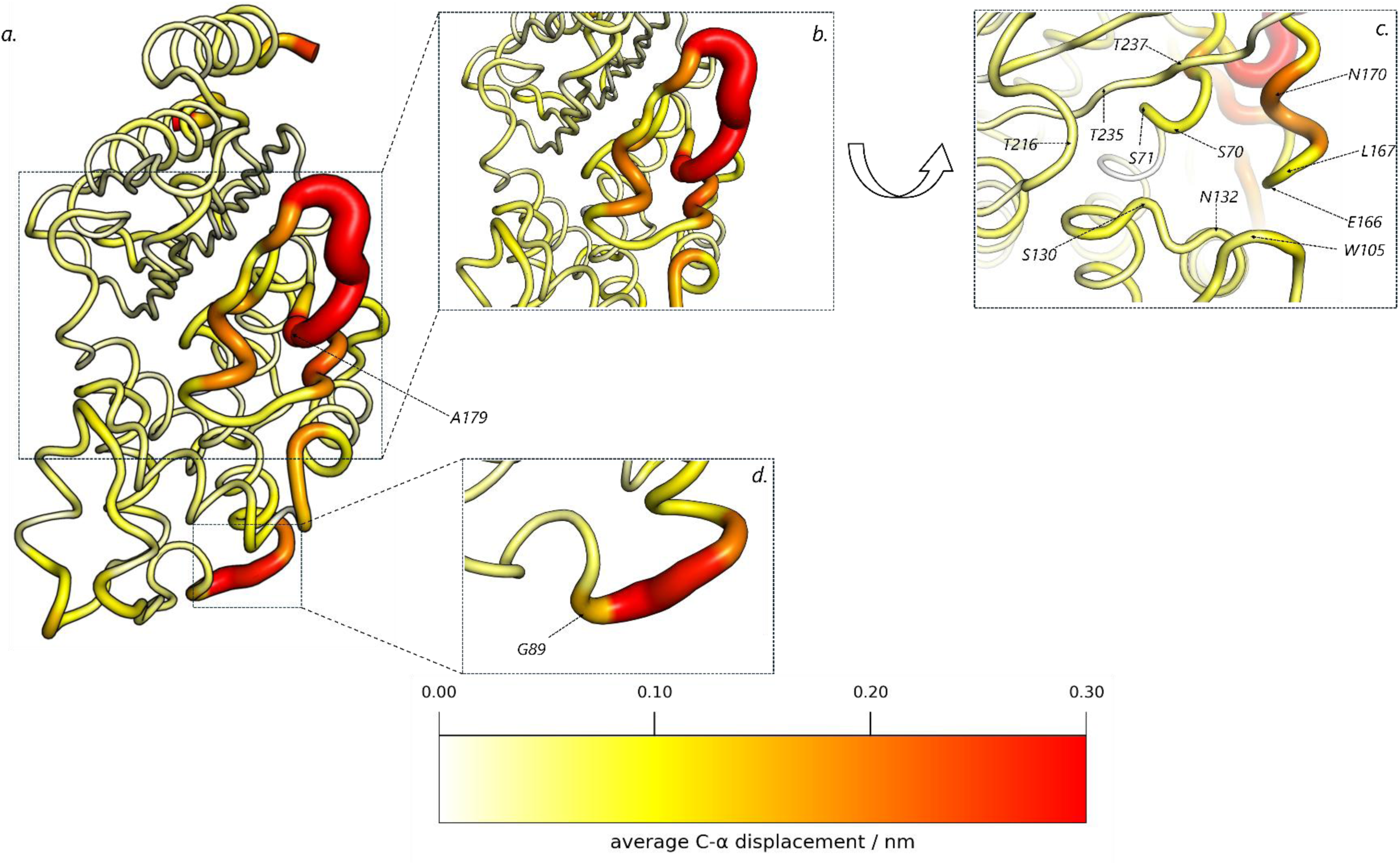
Structural response of KPC-2 aspartate to alanine mutation at position 179. ***a.*** Structural responses to D179A mutation at 1 ns of simulation time, with position 179 depicted as a sphere. Average Cα displacements are indicated at *t* =1 ns after the introduction of D179A. The displacements shown correspond to the norm of the average Cα displacement vectors after subtracting intrinsic protein fluctuations via the “null perturbation” analysis (2, 62), as detailed in Materials and Methods. The magnitude of displacements is colour-mapped from 0 to 0.3 nm according to the scale shown in the figure. The cartoon thickness also indicates the average responses to alanine mutation. D-NEMD responses show structural connectivity between position 179 and several functional regions in KPC-2, including the Ω-loop. ***b-d*** Detailed view of the average Cα displacements at the active site (***b***/***c***, rotated for a clearer view into the active site) and α2–β4 loop (residues 87–90) (***d***). Important residues such as E166, N170, and G89 are labelled and indicated with arrows.

The most pronounced changes upon alanine mutation are observed in the Ω-loop, mainly around the mutated residue 179. These responses quickly propagate to residues closer to the active site, specifically E166, L167 and N170 within the Ω-loop, all of which exhibit persistent displacements throughout the entire 1 ns (Figure S7d). Within the active site, residue N170 shows the largest average displacement, with an outward motion from the active site (Figure S3), which is likely to alter deacylation geometry and water positioning.

The mutation-induced structural changes observed in the Ω-loop have long-range effects on other key regions of the enzyme, namely the α3-α4 loop and hinge region (Figure 2). The α3-α4 loop contains the active site residue W105, which shapes the substrate-binding pocket through formation of aromatic stacking and hydrophobic contacts with various antibiotics, and has been implicated in substrate recognition and inhibitor resistance (27). In our simulations, consistent with the altered substrate profile of the D179A variant, W105 presented a mild increase in its average Cα displacement. Meanwhile, the hinge region, which includes T216, a residue known through experiments to be involved in binding the C3 carboxylate group of carbapenem substrates (25) also exhibited persistent structural responses.

Structural communication from position 179 also extends into the centre of the active site, likely propagated through the breaking of hydrogen bonds to residues 67 and 68. These contacts bridge the Ω-loop to the short loop preceding the catalytic centre, providing a pathway for structural rearrangements reaching to the S70 nucleophile, and beyond to other catalytically important residues. S130 — part of the serine-aspartate-asparagine (SDN) motif that participates in substrate anchoring, and in proton transfers following β-lactam bond cleavage (67); and N132, which helps orient and stabilise the substrate carboxylate and supports the local hydrogen-bond network around the catalytically important lysine 73, both show minor movements (0.06 and 0.05 nm respectively) (Figures 2 and S2). Comparable, mild displacements are detected in the region of the catalytic serine (S70), with S70 and S71 experiencing an average Cα displacement of 0.09 nm each. In the active-site β-sheet, T235 and T237, which respectively contribute to the carboxylate-binding pocket and the oxyanion hole (21), show small (0.03 and 0.05 nm) but statistically significant responses.

Notably, the α2–β4 loop (residues 87–90) also responds strongly to the introduced alanine substitution at residue 179, coinciding with heightened responses in the Ω-loop (Figure 2d). These results indicate that the Ω-loop and the α2–β4 loop are dynamically linked, with structural changes at position 179 influencing both sites. This relationship is consistent with our previous experimental observations (Beer *et al.* (2)) in which we reported that motions in the α2–β4 loop are coupled to conformational changes in the Ω-loop. The dynamical behaviour of the D179A variant described here therefore reinforces the existence of a functional and structural connection between these two loop regions.

Overall, the D-NEMD simulations show that position 179 exhibits robust, long-range coupling to the active site and α2–β4 loop, and also reveals the routes through which changes induced by mutation at this position are transmitted to these regions. The substitution of aspartate with alanine at position 179 strongly affects several key residues, including S70, S71, S130, N132, E166, L167, N170, T216, T235, T237; as well as the α2–β4 loop, all of which show statistically significant responses following the mutation. These findings highlight not only the intricate structural connectivity between position 179 and the KPC-2 catalytic centre, but also the influence of the D179A substitution on specific distal regions.

### Response to Alanine Mutation at Position 164

In experimental structures of KPC-2, residue R164 forms a salt bridge with D179, and a hydrogen bond with S171 (20) in the Ω-loop (Figure 1c). Disruption of these interactions by substituting R164 with alanine leads to patterns of signal propagation in D-NEMD simulations similar to those observed for the D179A variant, with the Ω-loop exhibiting the most pronounced responses (Figure 3).

**Figure 3:**
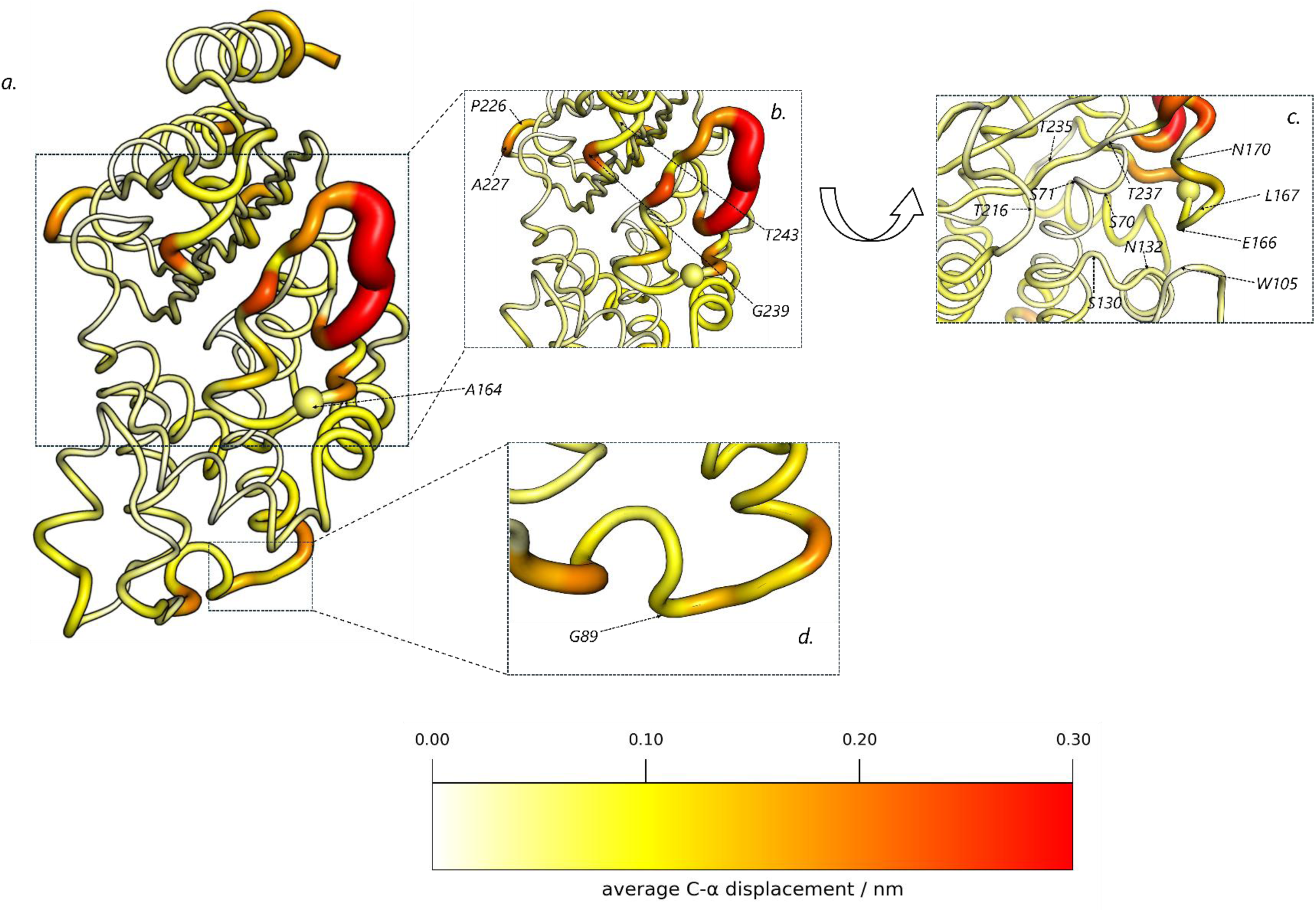
Responses from alanine mutation at position 164. ***a.*** Structural responses arising from R164A substitution at 1 ns of simulation time, with position 164 represented as a sphere. Mutation-induced changes are quantified by the average Cα displacements, determined after subtracting the intrinsic protein fluctuations via “null perturbation” analysis^2, 62^, as detailed in the Materials and Methods section. The responses are more localized than for D179A, probably due to residue 164 being more solvent-exposed. Despite this, D-NEMD responses show clear connection between position 164 and the Ω-loop (via residues 226, 227, 239-243 shown in ***b***), with active site residues (shown in ***c***, rotated for a clearer view into the active site), and with the α2–β4 loop (residues 87–90, shown in ***d***).

The active site residues E166, L167 and N170 in the Ω-loop all exhibit notable responses to alanine substitution at position 164 (displacements 0.05, 0.07 and 0.06 nm, respectively) but to a lesser (around 50% less) extent than for the D179A substitution (displacements 0.07, 0.10 and 0.20 nm, Figures 2-3). The Ω-loop responses are generally in similar directions (Figure S4).

Beyond the Ω-loop, the R164A substitution produces a distinct pattern of signal propagation compared with D179A. The β3–β4 loop (residues 239–243) plays a considerably larger role in transmitting structural changes, with displacement magnitudes of 0.22, 0.09, 0.07, 0.10 and 0.05 nm—substantially greater than the corresponding values for D179A (0.02, 0.07, 0.06, 0.07 and 0.03 nm) (Figure 3b). From this region, structural communication progresses into the enzyme core along the β-sheet scaffold, marking a muted but discernible communication path from the Ω-loop to the catalytic centre.

This pathway affects several central residues. The catalytic S70–S71 pair shows modest responses (0.03 and 0.01 nm), while residues S130 and N132 of the SDN motif undergo displacements of 0.03 and 0.04 nm (Figures 3, S6c). Further coupling extends to residues T235 (0.05 nm) and T237 (0.02 nm), that make interactions with the β-lactam carboxylate and oxyanion. Residue W105 (α3–α4 loop), a key gating residue for substrate binding, also responds along this route, consistent with signal propagation through the β-sheet rather than via a direct connection from the 239–243 loop.

A notable feature, unique to the simulations of the R164A substituted variant, is the appearance of measurable responses in the more distal residues 226 and 227 (Figure 3b), located at the end of the α11–β7 loop. These responses likely arise from signals propagated from the β3–β4 loop and transmitted through the central β-sheet (residues 234–238). Their involvement suggests that residues 226–227 may function as auxiliary nodes within an extended communication network, linking signals originating in the Ω-loop to peripheral regions, including the hinge loop. Consistent with this, residue T216 in the hinge region exhibits a displacement of 0.02 nm.

The R164A substitution also induces responses in the α2–β4 loop, although the magnitude of these is lower than is the case in D-NEMD simulations of the D179A variant (average displacement 0.12 vs 0.17 nm; Figure 3d). This attenuation likely reflects the more solvent-exposed position of R164, which weakens its coupling to the catalytic centre and consequently reduces the magnitude of signal transmission to the α3–β4 loop. The reduced displacements of S70–S71 further indicate that communication into this region is less extensive in the case of the R164A, compared to the D179A, substitution.

To summarise, in comparison with D179A, the R164A substitution elicits weaker structural changes in core residues, with an average decrease in response magnitude of approximately 52%. The lower impact of R164A is probably due to the more solvent-exposed location, which probably diminishes its influence on the core. Nevertheless, and despite the differences in behaviour between positions 179 and 164, D-NEMD clearly highlighted the pathways linking both residues that participate in the 164-179 salt bridge to the active site. This is consistent with experimental evidence that mutations at these sites affect enzyme activity and resistance profile, and their established roles in stabilising the Ω-loop and positioning catalytic residues (14, 19, 10, 29, 31, 65). Experimental data indicate that mutations of R164 have less effect on catalytic efficiency and substrate specificity than for D179 (14, 20, 65). The findings from the D-NEMD simulations described here are consistent with this conclusion: the R164A substitution induces weaker responses from active site residues than does D179A, i.e. the more buried position 179 is more strongly connected to the active site than the more solvent-exposed R164. Moreover, differences in the D-NEMD responses to alanine substitution at these two positions show each site to influence distinct distal regions, highlighting the nuanced and position-specific nature of structural perturbations within the enzyme.

### Response to Alanine Mutation at Position 220

The impact of a sequence change at position 220, in the hinge region, was also investigated by D-NEMD, due to the well-established role of this residue in the carbapenemase activity of KPC-2 (26). The arginine-to-alanine substitution (R220A) triggered notably rapid and widespread structural responses in the KPC-2 active site (Figure 4). Mutation-induced changes originating at position 220 rapidly propagated to the active site, resulting in large and permanent rearrangements in functionally important residues, including S70-S71, W105, T216, T235 and T237 (Figure S7a). These changes reached the active site more quickly than those observed for the R164A and D179 substitution, likely due to the closer proximity of position 220 to the enzyme’s catalytic centre. Displacements were observed for S70 (0.13 nm), S71 (0.10 nm), K73 (0.08 nm), S130 (0.08 nm), N132 (0.07 nm), W105 (0.15 nm), and T216 (0.13 nm), with these shifts persisting to 1 ns (Figure 4). Within the active site β-sheet backbone, T235 showed the largest response (0.17 nm) with a pronounced upward direction away from the active site (Figure S5), while T237 (0.12 nm) also exhibited a large average displacement, consistent with the disruption of the hydrogen bond network between this residue and position 220. As explained above, this hydrogen bond is essential for maintaining the correct orientation of T237 allowing for formation of hydrogen bonds to the C3 carboxylate group of carbapenem substrates (26). Compared to the previously tested positions (namely D179 and R164), sequence changes at position 220 induced structural responses of both larger magnitudes and a broader distribution, highlighting a particularly strong link between position 220 and the active site.

**Figure 4:**
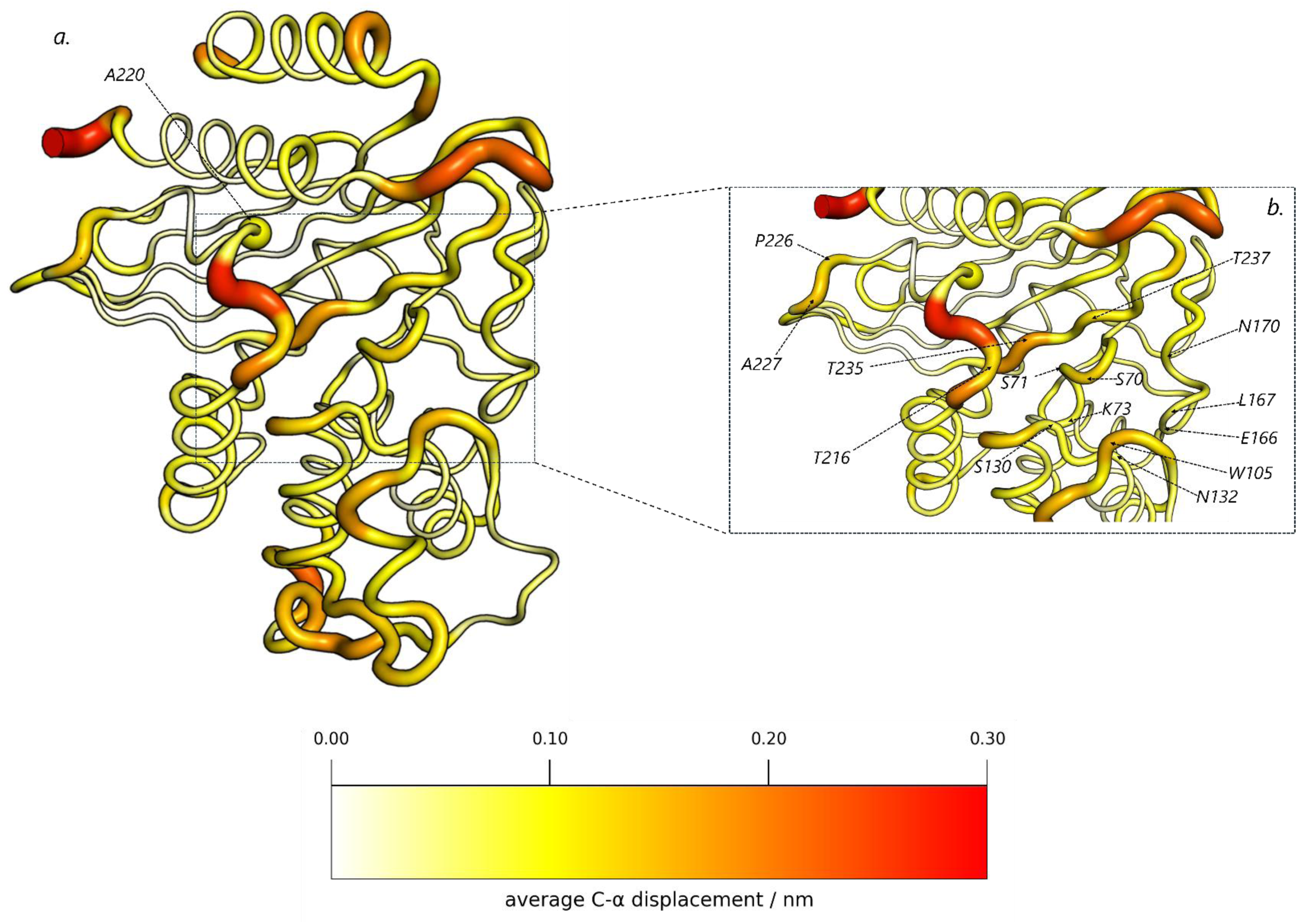
Response networks stemming from position 220. ***a.*** Structural responses arising from the R220A substitution at 1 ns of simulation time, with position 220 is represented as a sphere. Mutation-induced changes are quantified by the average Cα displacements, determined after removing the intrinsic protein fluctuations via the “null perturbation” analysis (2, 62), as detailed in the Materials and Methods section. The simulations point to position 220 having a key structural role within the enzyme, highlighted by an almost universal increase in the average Cα displacements throughout the enzyme, including the Cα atoms of the active site highlighted in ***b*.**

A key distinction between the behaviours of the D179A and R164A variants, and of the R220A substitution, lies in the response of the Ω-loop (Figures 2-4 and S6). When averaging the Cα displacements across all residues of the Ω-loop, R220A produces the smallest overall response, with a 75% reduction in mean Ω-loop displacement relative to the D179A, and a 72% reduction relative to the R164A, substitutions. This suggests a weakened coupling between the site of mutation (220) and this loop. Nevertheless, strong mutation-induced conformational changes still propagate (Figure 4) to active site residues E166, L167 and N170. Furthermore, the distal residues 226 and 227 (Figure 4b)— positioned at the start of the α11–β7 loop—exhibit moderate and consistent displacements in response to the R220A substitution, similar in magnitude to those observed in R164A. This behaviour further suggests that residues 226–227 may play a key role in the structural communication pathway between the Ω-loop and the hinge region.

Overall, substitution at position 220 induced rapid and profound structural responses from active site residues in the core of the enzyme, propagated through the hinge region. Although the Ω-loop showed reduced displacements in response to the R220A, compared to the D179A and R164A substitutions, conformational changes still reached it via the 239–241/242 loop. These findings suggest that position 220 plays a uniquely influential role in KPC-2 activity, consistent with experimental data showing an 80% reduction in carbapenemase activity upon mutation to alanine. The proximity of residue R220 to the catalytic centre may explain the faster and broader propagation of structural responses observed in the D-NEMD simulations.

### Effects of Substitutions at Position 276

Up to this point, our D-NEMD analysis has focused exclusively on positions where mutations are known to impact KPC-2 activity, with their individual communication networks to the active site identified. However, to rigorously evaluate the ability of the D-NEMD methodology to distinguish between functionally relevant positions (i.e., those at which substitution alters enzyme activity) and those where substitution exerts minimal or no effect, we investigated the effects of mutation to alanine of residue 276, for which experiments indicate that mutation does not affect catalytic activity (26). Residue E276 was selected because it is not only positioned near the active site, but also participates in hydrogen bonding interactions with residues R220 and T237, suggesting the potential for functional relevance. However, somewhat unexpectedly, kinetic data indicate that substitutions at E276 do not significantly alter the enzyme’s activity profile towards a range of β-lactam antibiotics (26). This might imply that residue 276 has only a weak, or lacks, connection to the catalytic centre. Including the E276A substitution then provides a control perturbation, analysis of which will enable us to evaluate the ability of D-NEMD to correctly identify substitutions of limited functional impact and distinguish the effects of perturbations at such positions from those at functionally important positions (i.e. that modulate activity).

Consistent with the results of experiments (26), the structural responses triggered by the E276A substitution after 1 ns of D-NEMD simulation were minimal and limited compared to those induced by alanine substitutions at positions 164, 179, or 220 (Figures 5 and S7-S8). Overall, minor conformational changes were observed in the active site (Figure 5), with only T237 and T216 displaying persistent displacements (0.07 nm each) (Figure S8). The response of T237 reflects the loss of its hydrogen bond connection to E276 via R220, while the shift in T216 likely arises from indirect effects due to its proximity to R220. No broader networks involving active site residues were observed following exchange of E276 to alanine, reinforcing the limited structural impact of the substitution (Figure 5). These results confirm position 276 as functionally unimportant, with very weak and limited links to the KPC-2 active site, validating its use as a control site.

**Figure 5:**
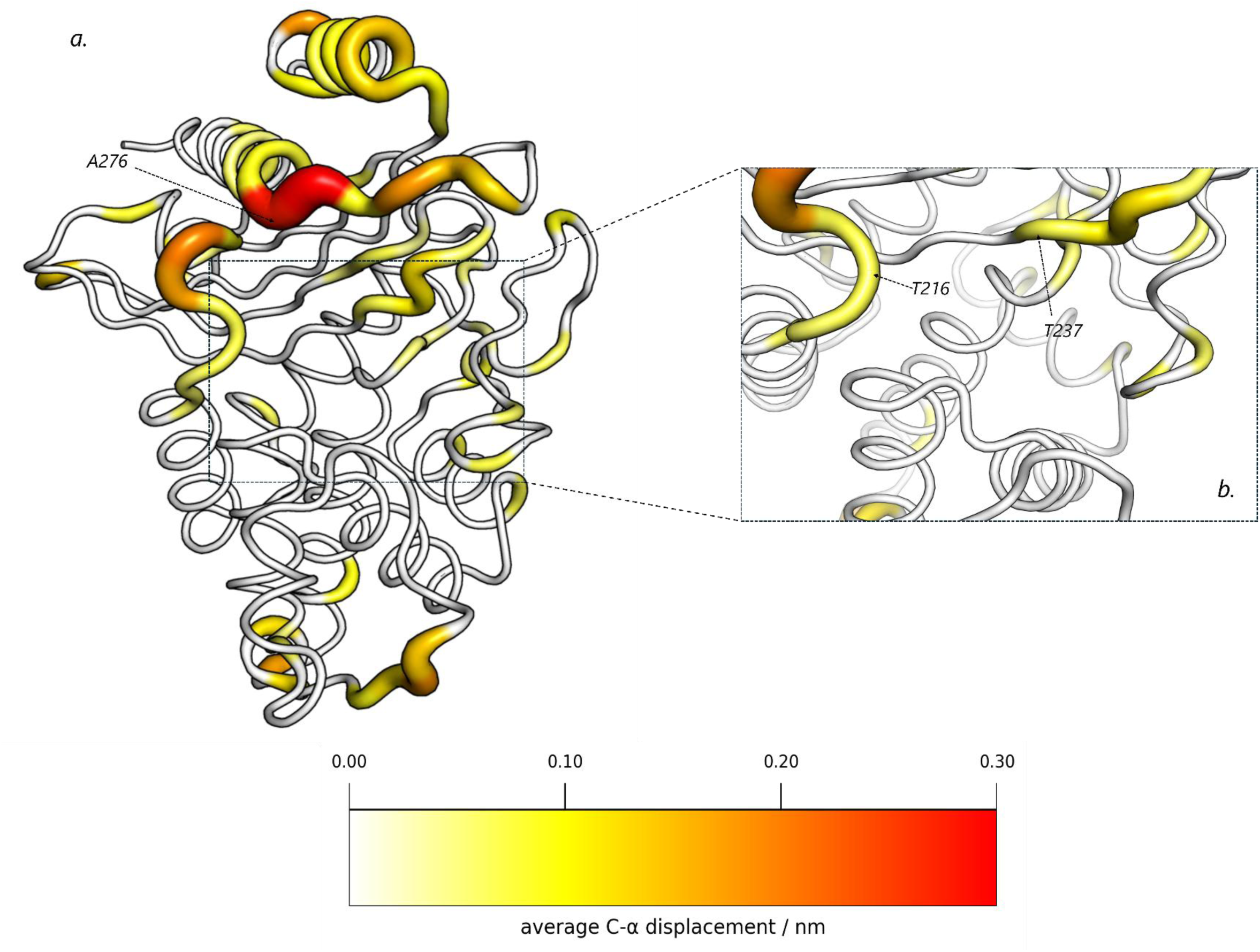
Response networks stemming from position 276. ***a.*** Structural responses arising from the E276A substitution, with position 276 represented as a sphere. Mutation-induced changes are quantified by the average Cα displacements, determined after removing the intrinsic protein fluctuations via the “null perturbation” analysis (2, 62), as detailed in the Materials and Methods section. D-NEMD responses reveal limited signal propagation to the active site or other key regions of the enzyme, following E276A substitution. Structural changes induced by the mutation remain largely confined to the local environment, with increased displacement observed at T237 due to disruption of the E276–T237 hydrogen bond. A response is also detected at R220, which transmits it to active site residue T216, highlighted in which shows a sustained increase in displacement. Highlighted in ***b.***, T237 and T216 are the only active site residues displaying prolonged responses. The response patterns captured by D-NEMD indicate that sequence changes at position 276 do not generally affect the dynamic behaviour of the enzyme’s active site. This is in line with experimental findings showing that the E276A substitution does not significantly alter the spectrum of activity of KPC-2 (26).

In summary, the largest structural responses following alanine substitution at E276 were confined to residues in direct contact with or in close proximity to the site of mutation, whereas the perturbation exerts minimal effects upon the KPC-2 active site. This limited propagation provides a structural basis for the experimental findings showing that mutations at residue E276 do not significantly affect the activity of KPC-2 towards different β-lactam antibiotics (26). These results also demonstrate the sensitivity of the D-NEMD methodology, in capturing subtle differences in internal communication networks, effectively distinguishing between sites that are functionally relevant, and those that are not.

Beyond the specific insights obtained for KPC-2, this work suggests that D-NEMD simulations of alanine substitution provide a broadly applicable and efficient strategy for probing communication in proteins and potentially identifying functionally important sites. This method enables a systematic assessment of how each position influences functional regions. This approach requires only a standard equilibrium MD dataset; once generated, the same ensemble can be reused to perform a relatively high throughput, *in silico* alanine scan without the need for mutation-specific equilibration. Hundreds of mutations could potentially be evaluated at relatively little additional computational cost, allowing dynamic effects of sequence changes to be identified to guide experiment. This method should aid in prioritising distal sites for experimental characterisation and help focus mutagenesis and engineering campaigns. This strategy can be applied to any protein for which equilibrium MD simulations are available.

## Conclusions

In this work, we present a novel application of D-NEMD simulations, in which we use (instantaneous) alanine substitutions to probe how local structural changes propagate through an enzyme and influence its dynamics. The perturbations induced by mutation of specific amino acid residues to alanine trigger dynamical responses and reveal the internal communication networks operating within the protein. The mutation-induced responses identify functionally relevant sequence positions, based on their ability to transmit structural signals to the active site, and other key regions, of the enzyme under investigation. Using the well-characterised class A β-lactamase KPC-2 as a model system, we demonstrate that D-NEMD distinguishes functional (i.e. those that alter enzymatic activity) and functionally unimportant positions, based on whether their substitution initiates long-range, structured responses that reach the active site.

Alanine substitutions at functionally significant positions in KPC-2, such as residues 179, 164 and 220, disrupt local stabilising interactions and generate persistent, time-resolved responses across the Ω-loop and extending to catalytically important residues, including E166 and N170. These effects extend far beyond the mutated positions, into conserved active site elements and neighbouring secondary structures, consistent with long-range communication between the sites of mutation and the active site. In contrast, substitutions at residues not linked to function, such as position 276, do not elicit major responses within the KPC-2 active site, underscoring their limited ability to modulate active site behaviour. These finding clearly show the utility of D-NEMD in identifying sites with functional connectivity, demonstrating how the method can effectively map allosteric networks linking distal sites to the active site, in the case of KPC-2 helping to explain how substitutions associated with changes in antimicrobial susceptibility of producer organisms exert their functional effects.

By tracking the direction and extent of displacements induced by alanine substitution, we identify structural pathways that connect sites of mutation to different functionally relevant regions in KPC-2. These new insights help explain how distal mutations can influence catalysis and provide a mechanistic basis for interpreting evolutionary changes in β-lactamases. With >245 KPC-type enzymes reported as of January 2025 (12), and new mutations continuing to emerge, many of these variants remain uncharacterized. New variants often emerge under therapeutic pressure and can significantly alter the enzyme’s susceptibility profile, complicating treatment strategies and clinical decision-making. This underscores the urgent need for reliable predictive tools to assess the functional impact of novel mutations. As shown here, D-NEMD (using alanine substitution as the external trigger) offers a powerful computational approach to quickly evaluate whether sequence changes are likely to influence enzymatic activity via long-range allosteric communication, helping prioritize experimental efforts to characterise variants of greatest clinical relevance, and identify functionally connected residues as potential targets for inhibitor design.

In summary, this study establishes alanine-substitution D-NEMD as a powerful and generalisable tool for probing the dynamic internal communication networks that govern protein function. By capturing the evolution of mutation-induced structural changes, D-NEMD goes beyond, and complements, equilibrium MD methods and offers a relatively rapid, structure-based strategy for predicting residues involved in allosteric networks, and to predict the impact of sequence changes. Its broad applicability promises to make it a valuable tool for probing allosteric communication in proteins, guiding experiments and informing protein design and engineering.

## Author Contributions

AJM, ASFO, JS devised the experiments with input from BB and MB. BB implemented the D-NEMD simulation setup and carried out the simulations under the supervision from MB and ASFO. BB analysed the simulations with supervision from MB and ASFO and input from DP. The manuscript was written by BB with input and edits from all authors. BB and DP are supervised in their PhD work by AJM.

## Acknowledgements

This work forms part of a project that received funding from the European Research Council under the European Horizon 2020 research and innovation program (PREDACTED Advanced Grant Agreement no. 101021207) awarded to AJM and JS. BB, MB and DP are supported by the European Research Council grant PREDACTED Advanced Grant Agreement no. 101021207. ASFO was supported at the University of Bristol by a BBSRC Discovery Fellowship ([BB/X009831/1]). ASFO thanks the University of Bristol “Heather Corrie Impact Fund” for support. This work used computational resources provided by the University of Bristol Advanced Computing Research Centre (ACRC). We thank EPSRC for providing ARCHER2 HPC time via HECBioSim (hecbiosim.ac.uk).

## Data and code availability

All D-NEMD simulation data, including input and trajectory files, will be made openly accessible via the University of Bristol data repository, data.bris, upon manuscript acceptance.

