## Supporting Information for "A dynamical-nonequilibrium molecular dynamics (D-NEMD) alanine scanning approach for identifying allosteric positions in proteins"

### Supporting Figures

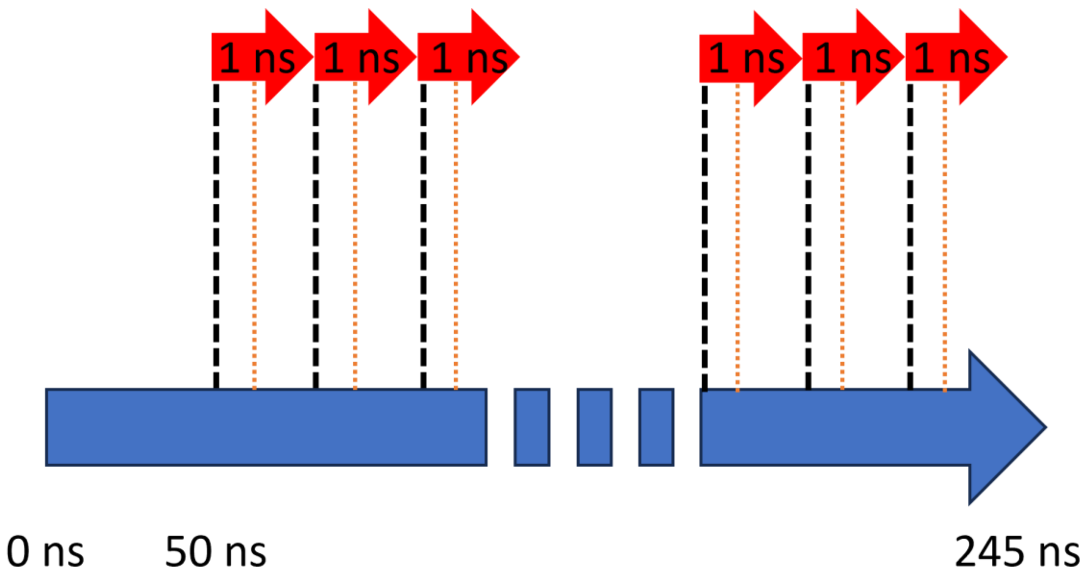


**Figure S1: Graphical representation of D-NEMD workflow.** The large blue arrow represents one of 5, 250 ns long, equilibrium MD simulation of KPC-2. Dashed black lines represent system snapshots taken every 5 ns along the equilibrium trajectory. From each of the five equilibrium MD repeats, 40 frames were extracted, resulting in a total of 200 conformations used as starting points for D-NEMD. In each snapshot, the target residue was substituted with alanine, and simulated for 1 ns (the resulting nonequilibrium simulations are illustrated by red arrows). The evolution of KPC-2 response to alanine mutation was extracted using the Kubo-Onsager relation[57]. Comparisons between equilibrium and nonequilibrium trajectories at equivalent timepoints were (indicated by orange lines) were averaged across all repeats to obtain statistically significant dynamical responses.

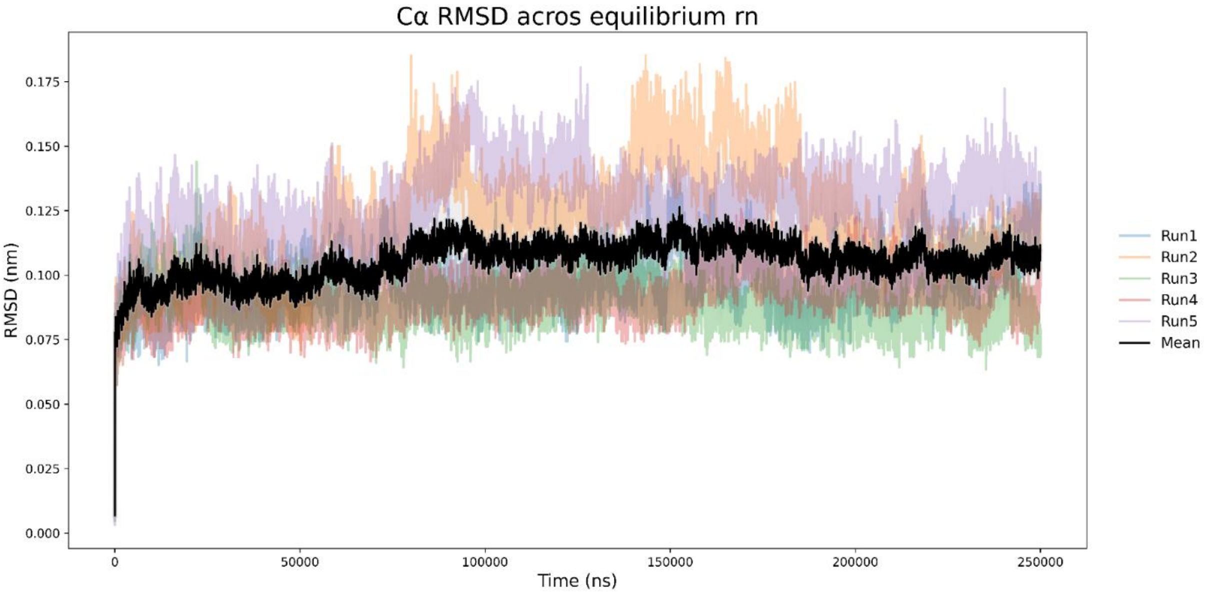


**Figure S2: Cα root mean square deviation (RMSD) of equilibrium simulations.** Cα RMSD of five independent equilibrium simulations of KPC-2 (PDBID: 6D16). The first and last five residues were excluded from the analysis to avoid contributions from highly flexible termini. Individual simulation traces are shown in semi-transparent lines, with the mean RMSD indicated by a solid black line. RMSD values were calculated relative to the initial structure following least-squares fitting of the included residues.

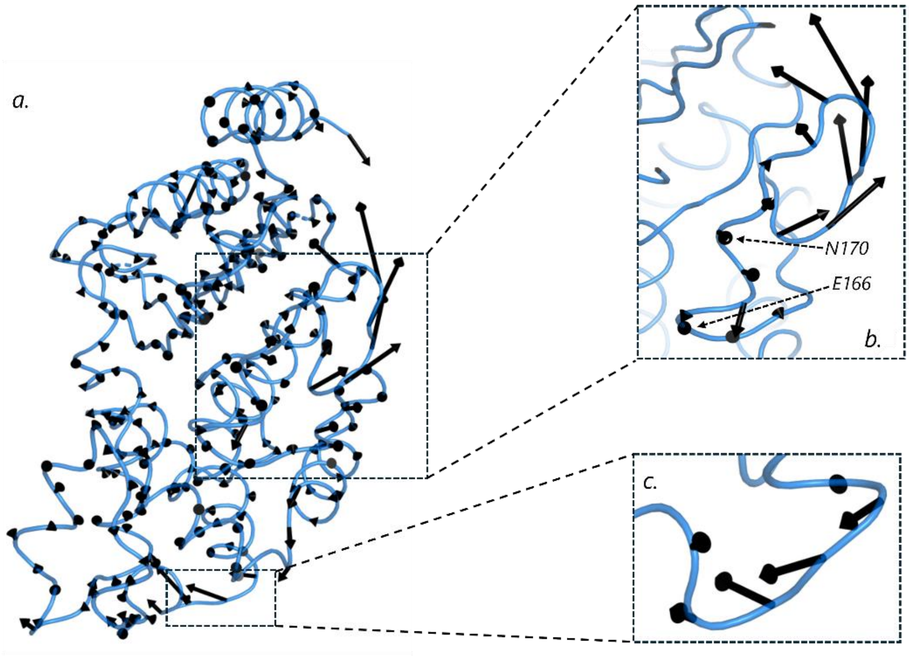


**Figure S3: Directionality of KPC-2 structural response to alanine substitution at position 179 *a.*** Average Cα displacement vectors at 1 ns following alanine mutation at D179 ***b.*** Detailed view of the Ω-loop responses, where the most significant response vectors are observed, including active site residues E166 and N170. ***c.*** Close-up view the concurrent responses in the α2–β4 loop (residues 87–90). The black arrows correspond to the average Cα displacement vectors, determined between the equilibrium (D179) and nonequilibrium (D179A) trajectories over the 200 replicas and after subtracting intrinsic protein motions via the "null perturbation" analysis with only statistically significant responses plotted, as detailed in the Materials and Methods. Vectors with a length 0.15 nm are displayed with a scale-up factor of 2.

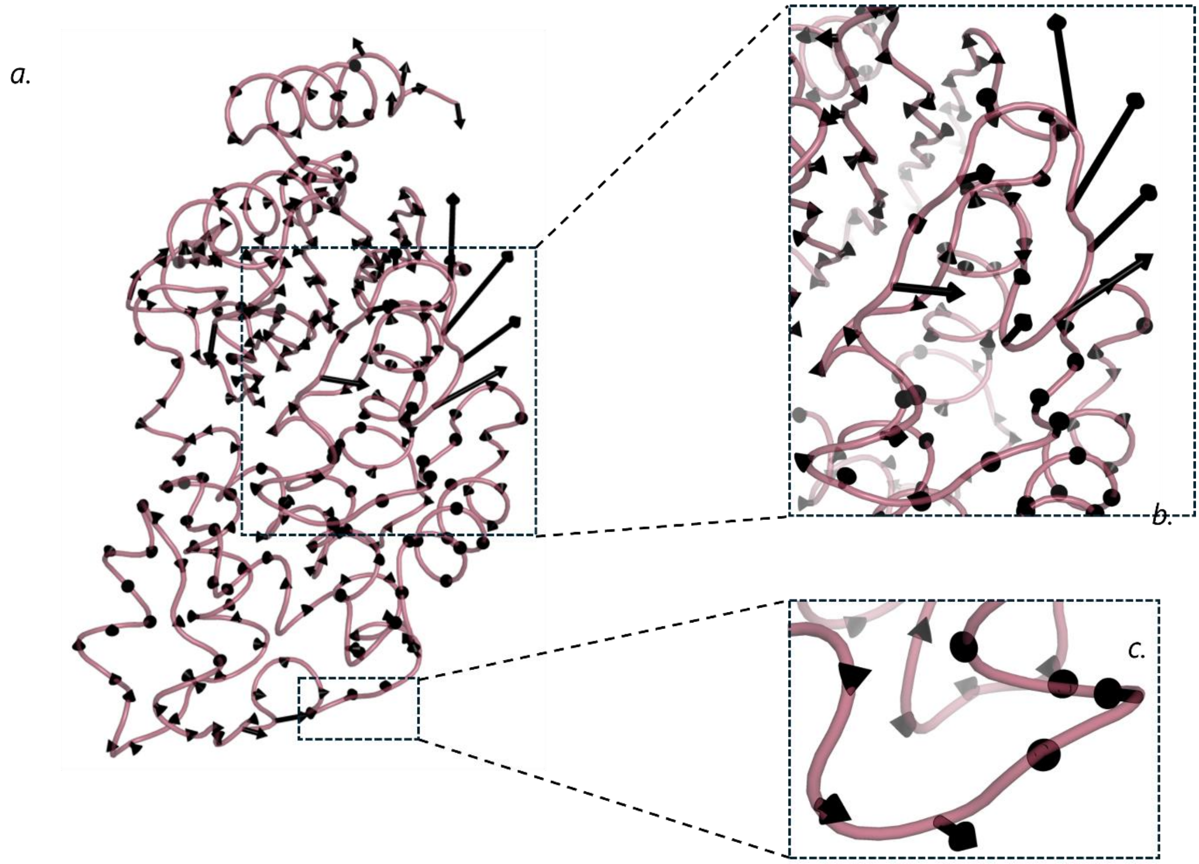


**Figure S4 Directionality of KPC-2 structural response to alanine substitution at position 164.** ***a.*** The figure shows the structural responses initiated by the R164A substitution at 1 ns of simulation time. Mutation-induced changes are quantified by the average vector of Cα atoms, determined after removing the intrinsic protein fluctuations via the "null perturbation" analysis with only statistically significant responses plotted, as detailed in the Materials and Methods section. ***b.*** Ω-loop, where the largest responses are observed, including active site residues E166 and N170. ***c.*** Responses of the α2–β4 loop (residues 87–90), although much less pronounced than in D179A. Vectors with a length 0.15 nm are displayed with a scale-up factor of 2.

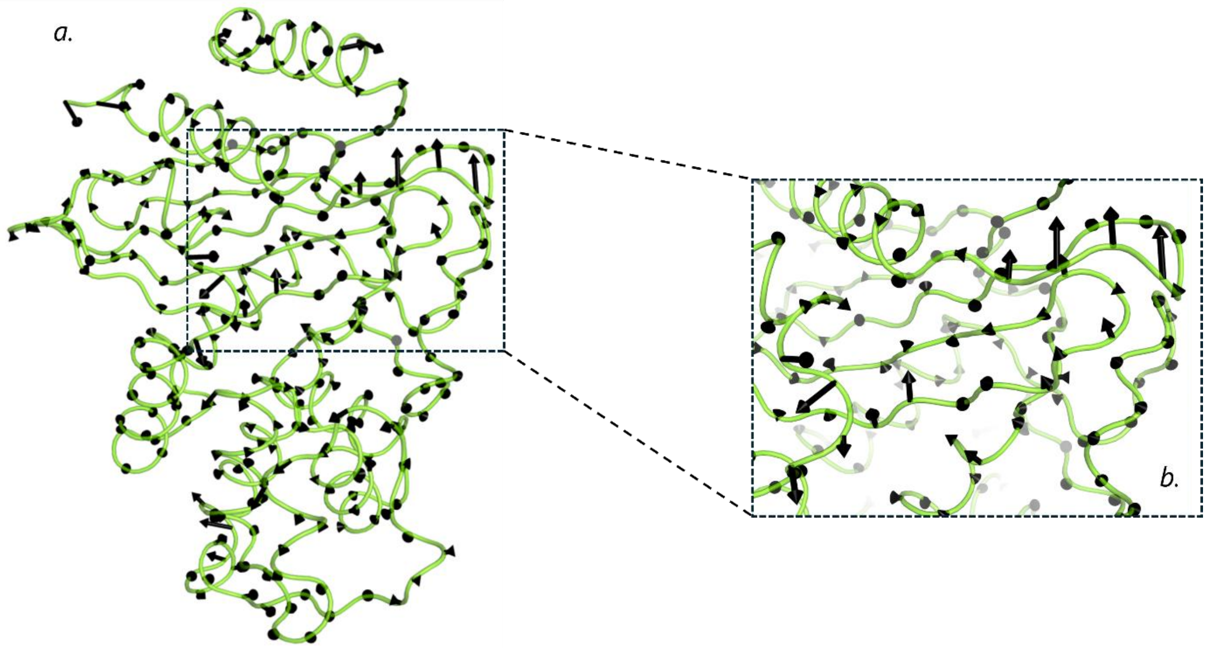


**Figure S5: Directionality of KPC-2 structural response to alanine substitution at position 220. *a.*** The figure shows the structural responses initiated by the D179A substitution at 1 ns of simulation time. Mutation-induced changes are quantified by the average vector of Cα atoms, determined after removing the intrinsic protein fluctuations via the "null perturbation" analysis with only statistically significant responses plotted, as detailed in the Materials and Methods section. ***b.*** Highlighting the hinge region (residues 213-220), the β-sheet backbone of the active site (residues 234-237) and residues 270-276, where the most significant response vectors are observed, including active site residues T216, T235 and T237. Vectors with a length 0.15 nm are displayed with a scale-up factor of 2.

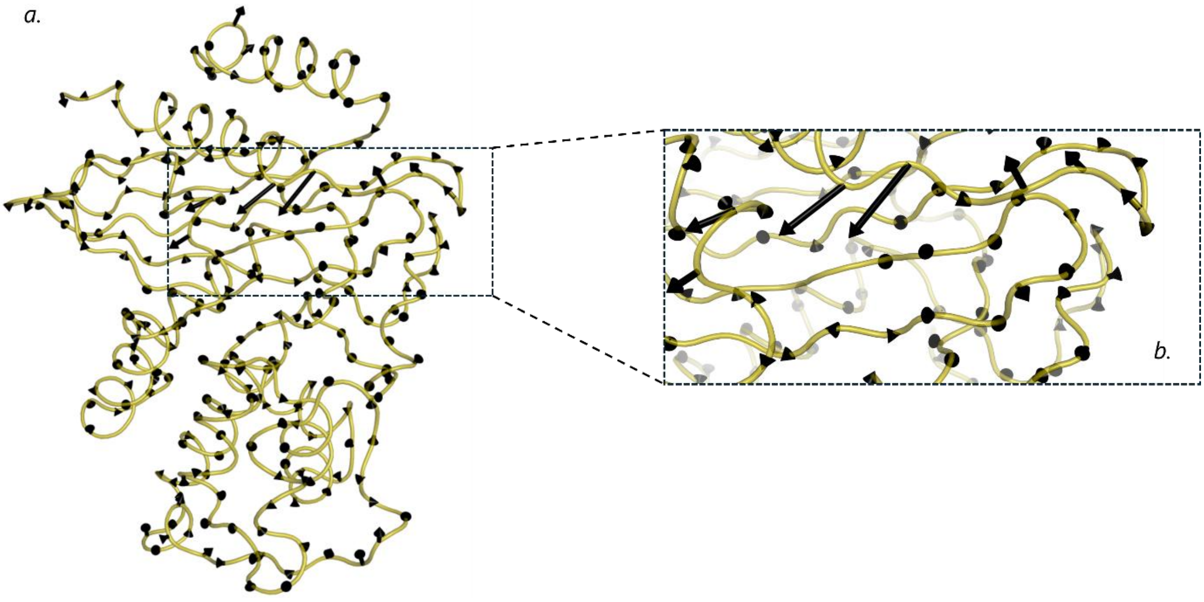


**Figure S6: Directionality of KPC-2 structural response to alanine substitution at position 276.** ***a.*** The figure shows the structural responses initiated by the D179A substitution at 1 ns of simulation time. Mutation-induced changes are quantified by the average vector of Cα atoms, determined after removing the intrinsic protein fluctuations via the "null perturbation" analysis with only statistically significant responses plotted, as detailed in the Materials and Methods section. ***b.*** Highlighting the hinge region (residues 213-220), the β-sheet backbone of the active site (residues 234-237) and residues 270-276, where the most significant response vectors are observed. Active site residues are notably not undergoing pronounced responses. Vectors with a length 0.15 nm are displayed with a scale-up factor of 2.

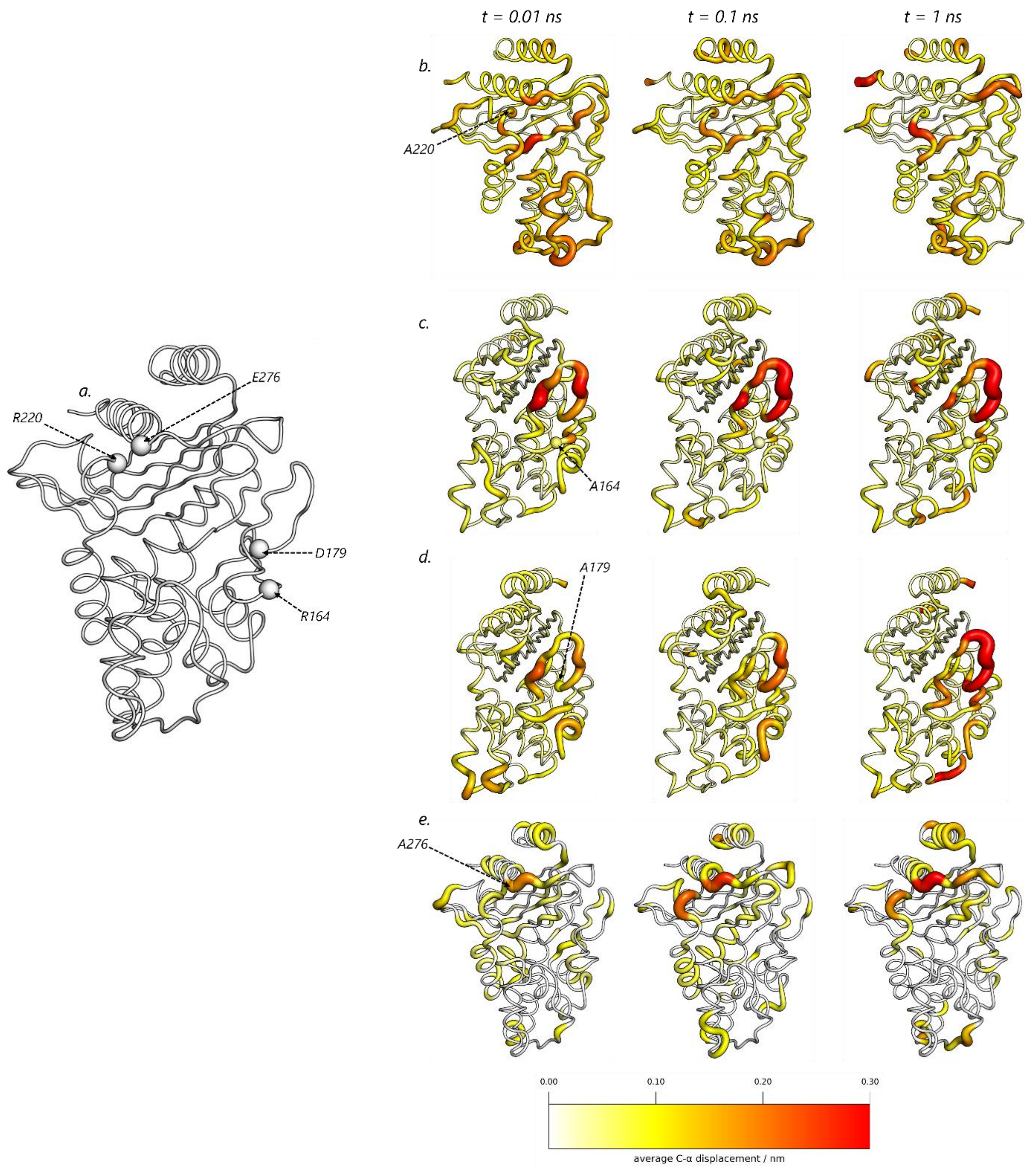
t = 0.01 ns
t = 0.1 ns
t = 1 ns
a.
R220
E276
D179
R164
b.
A220
c.
A164
d.
A179
e.
A276
0.00
0.10
0.20
0.30
average C-α displacement / nm

**Figure S7: Structural response of KPC-2 to alanine substitutions. *a*.** Location of the four sites selected for alanine substitution using D-NEMD. The position of the R164, D179, R220, and E276 is represented with spheres. ***b-e*.** Average Cα displacements at *t = 0.01, 0.10* and *1.00 ns* after alanine substitituion at position R220 (***b***), R164 (***c***), D179 (***d***), and E276 (***e***). The displacements shown correspond to the norm of the average Cα displacement vectors after removing the intrinsic protein fluctuations via the "null perturbation" analysis, as detailed in the Materials and Methods. Both the cartoon thickness and structure colours (mapped from 0 to 0.3 nm according to the scale) indicate the average responses to alanine mutation.

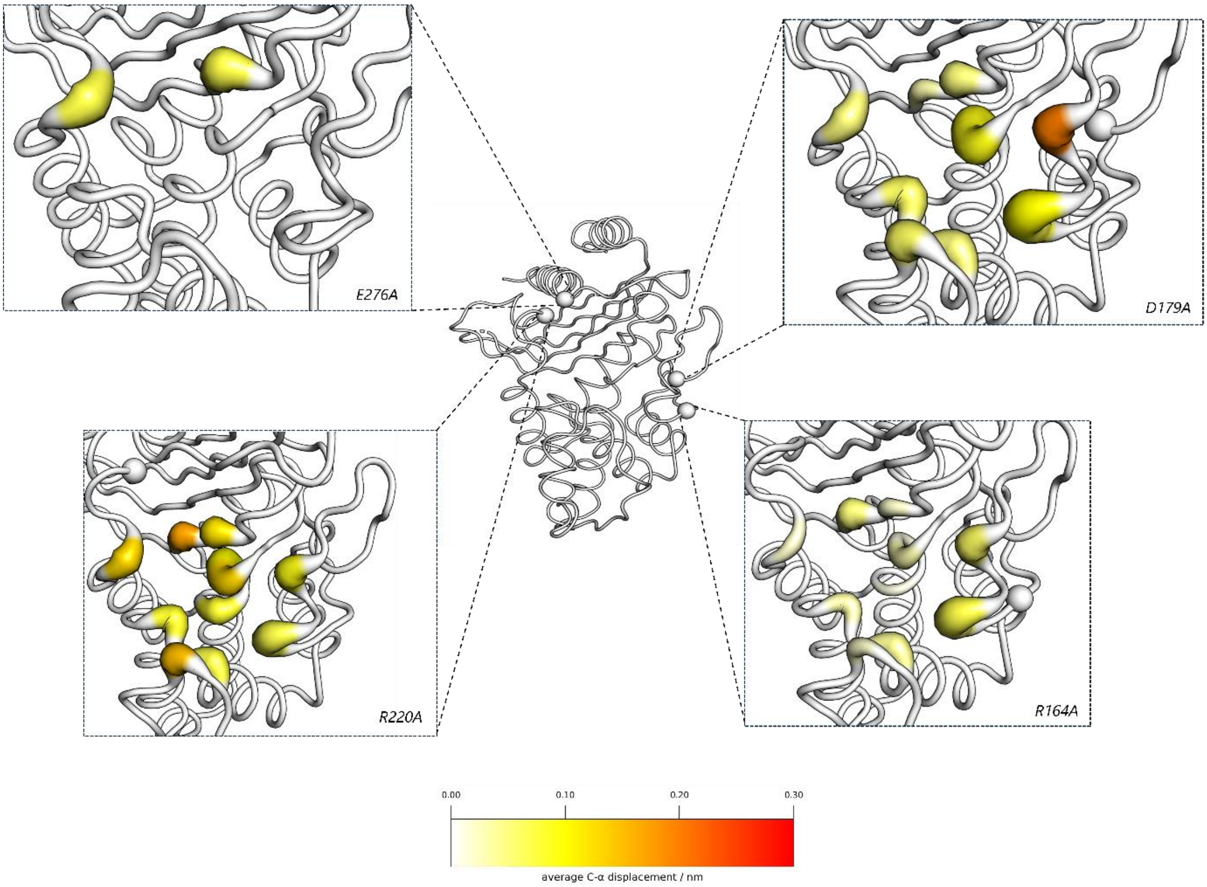


**Figure S8: Structural responses of KPC-2 active site residues to alanine substitutions.** The figure shows the structural changes initiated by alanine substitution in the active site residues of KPC-2 at *t =1000 ps* . The white spheres represent the Cα atom of the alanine-mutated residues, namely E276A (top left), D179A (top right), R220A (bottom left) and R164A (botton right). The displacements shown correspond to the norm of the average Cα displacement vectors after removing the intrinsic protein fluctuations via the "null perturbation" analysis, as detailed in the Materials and Methods. Both the cartoon thickness and structure colours (mapped from 0 to 0.3 nm according to the scale) indicate the average responses to alanine mutation. Please note the clear difference in the extent of structural responses propagated to the active site across variants, with the functional substitutions, D179A, R164A and R220A, inducing larger, widespread and persistent changes in active site residues, indicating a clear and strong structural connectivity between the mutated positions and the active site. In contrast, E276A elicits limited and more subtle structural responses within the active site, primarily affecting T237 and T216 through hydrogen bond rearragment, with no broader engagement of other active site residues.

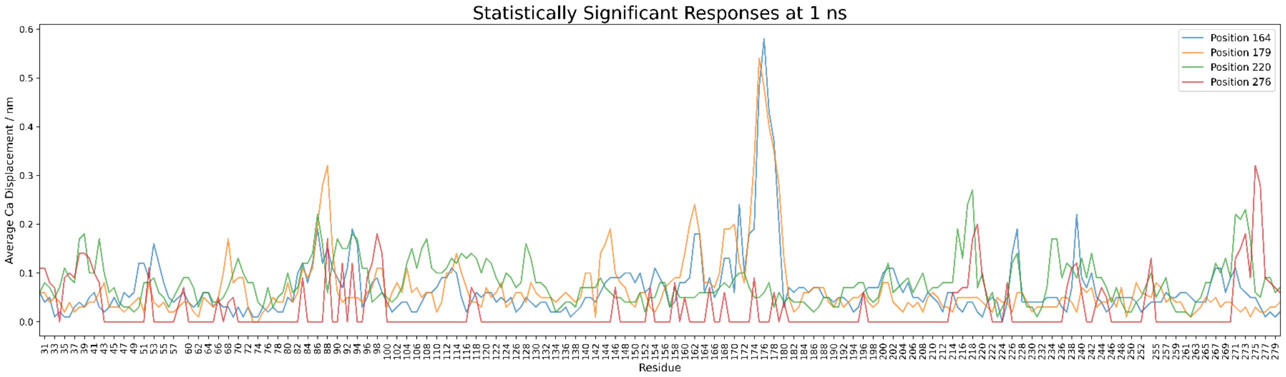


**Figure S9: 2D Plot of structural responses of all mutants.** Two-dimensional representation of residue-specific structural responses for all mutants at 1 ns of simulation time. The displacements shown correspond to the norm of the average Cα displacement vectors after removing the intrinsic protein fluctuations via the "null perturbation" analysis, as detailed in the Materials and Methods. This plot enables comparison of the magnitude and distribution of conformational changes across different mutant simulations.
